# Neurosphere culture derived from aged hippocampal dentate gyrus

**DOI:** 10.1101/2024.03.16.585365

**Authors:** Olga Vafaeva, Poommaree Namchaiw, Karl Murray, Elva Diaz, Hwai-Jong Cheng

**Author notes:** Correspondence: Elva Díaz,; or Hwai-Jong Cheng.

## Abstract

The neurosphere assay is the gold standard for determining proliferative and differentiation potential of neural progenitor cells (NPCs) in neurogenesis studies^1–3^. While several *in vitro* assays have been developed to model the process of neurogenesis, they have predominantly used embryonic and early postnatal NPCs derived from the dentate gyrus (DG). A limitation of these approaches is that they do not provide insight into adult-born NPCs, which are modeled to affect hippocampal function and diseases later in life. Here, we show a novel free-floating neurosphere culture system using NPCs isolated from the DG of mature adult and aged mice.

The protocol outlines detailed steps on the isolation, propagation, and maintenance of neurospheres from adult and aged (>12 months old) mouse brain and how to differentiate cultured neurospheres into neurons and astrocytes. Culturing adult and aged NPCs provides an important *in vitro* model to (1) investigate cellular and molecular properties of this unique cell population and (2) expand the understanding of plasticity in the adult and aging brain. This protocol requires ∼2 hours to complete dissection, dissociation and culture plating, while differentiation to neuronal and astrocytic lineages takes 9 days.

By focusing on neurospheres obtained from animals at later ages this model facilitates investigation of important biological questions related to development and differentiation of hippocampal neurons generated throughout adult life.

## Introduction

Adult neurogenesis refers to generation of newborn neurons from resident stem and neural progenitor cells (NPCs) in mature (adult) animals. In mammalian brain, two neurogenic niches remain active throughout life, namely the subventricular zone (SVZ) of the lateral ventricle and the subgranular zone (SGZ) of the dentate gyrus (DG) of the hippocampus^4–7^. In the SGZ, these cells generate granule cell (GC) neurons in the DG of the hippocampus which play an important role in memory and learning processes^8–11^. While adult neurogenesis persists throughout life the proliferative capacity of NPCs declines with age^12,13^. Understanding how adult-born GCs differentiate and integrate into the hippocampal circuitry later in life requires a model that allows for high resolution analysis at the single cell throughout the lifespan.

The neurosphere assay (NSA) is a widely used *in vitro* technique to isolate and determine the proliferative and differentiation potential of stem cells and NPCs in rodent models ^2,3,14^. It enables investigation of this complex process by recapitulating *in vivo* events, while also reducing known and unknown confounding variables present *in vivo*^1^. Importantly, this technique allows for expansion of the number of dividing cells and establishment of a homogeneous reservoir that can be cryopreserved for future experiments.

These features make the NSA particularly useful for investigating cellular and molecular drivers of age-related reduction of adult neurogenesis. Since the population of NPCs in the adult brain is low and further decreases with age, this poses a technical challenge for detection and quantification of neurogenesis events *in vivo* such as cell growth, differentiation, integration and death. Usually, such studies rely on cell labeling and lineage tracking approaches followed by labor-intensive and time-consuming histological methods of tissue fixation, sectioning, staining and microscopic analysis. Morphological development and distribution of newly generated cells can also be approached with *in vivo* imaging methods such as miniscope implantation^15,16^. This approach allows for longitudinal studies of adult neurogenesis. However, it requires expensive specialized equipment and extensive training in surgical and post operative animal care procedures. The NSA, as described in this protocol, provides a useful model to investigate cellular and physiological mechanisms that control different steps of neurogenesis such as proliferation, cell death, fate choice and synapse formation and maturation *in vitro*. It allows for production of increased quantities of NPCs which is important in the context of decreased neurogenesis later in life. These cells can be cryopreserved allowing for multiple downstream assays to be performed on cells derived from the same animal and at coordinated lineage stages which can reduce variation in experimental measures. This protocol provides a valuable technique to test hypothesis on mechanisms of cell autonomous regulation of NPC survival, proliferation and differentiation in the adult and aged brain.

### Development of the protocol

Our laboratory has a longstanding interest in understanding how molecular identity, cellular differentiation, and synaptic integration of hippocampal neuronal progenitors are altered during the aging process. Previously, we described the dynamics of synaptic spatiotemporal development and the cellular process by which adult born neurons functionally integrate over time changes throughout the lifespan^17,18^. We observed a differential mechanism of maturation and synaptic integration of adult-born neurons in aged brain that could result from changes in intrinsic cell autonomous properties.

Herein, we describe the protocol for culturing NPCs from adult and aged animals established in order to study differentiation and integration of adult-born NPCs. To isolate and establish neurosphere cultures from mice >12 months old, it is necessary to account for lower numbers of actively dividing neural precursor cells and less efficient enzymatic degradation of tissue of adult and aged brain. To overcome these issues, we developed and integrated a novel step allowing the initial dissociated cells to proliferate on a coated plate so the rare progenitor cells to reach confluence before replating as free floating neurosphere cultures. We also found that the proprietary enzymatic cocktail provided in the Neural Dissociation kit for Postnatal neurons from Miltenyi Biotec provides the best dissociation and cell viability outcome, allowing culturing of neurospheres from mice up to 17-months old. It also, potentially provides, more consistent and more reproducible results between different users. The combination of longer culturing times after plating and more efficient papain-based tissue dissociation allowed us to develop a reliable protocol to grow, expand, and differentiate neurospheres from aged mice.

An important consideration is that this NSA protocol might not be effective in isolating and propagating quiescent stem cells. Quiescent stem cells are in the G_0_ phase of the cell cycle and therefore not actively dividing whereas this NSA primarily selects for cells that are undergoing proliferation, allowing for their expansion *in vitro*.

### Applications of the method

Neural precursor cells are identified in adult brains of numerous species including fruit flies ^19^, teleost fish ^20^, birds ^21^ and mammals^4,5,22^. Owing to their relative rarity in brain tissues these cells are challenging to study *in vivo.* The NSA described herein allows isolation of NPCs from adult and aged mouse brain, expansion of dividing cells and generation of a homogeneous reservoir of cells that can be cryopreserved for future experiments. The ability of these cell lines for self-renewal and generation of differentiated progeny demonstrates that the NSA represents a sound approach for isolating and propagating NPCs from DG across all ages. Multiple neurosphere cultures can be established from mice of the same age to serve as biological replicates and increase rigor of experimental applications. Moreover, neurosphere cultures generated from mice of both sexes provides a mechanism to investigate sex as a biological variable.

The NSA has been successfully used to understand mechanisms of cell proliferation, self-renewal and differentiation in NPCs generated postnatally. Combined with animal models of disease or transgenic animals this technique could be used to study how different steps of neurogenesis are affected in pathological conditions. In particular, most mouse models of neurodegeneration studies don’t show pathology until animals are relatively old. The ability to isolate NPCs from aged mice in the current protocol will greatly facilitate such analysis.

Adult NPCs derived as neurospheres can also be used for molecular analysis of RNA and/or protein expression. Since the NSA allows for the isolation and expansion of NPCs *in vitro* relatively rapidly, it enables higher throughput and shorter experimental turn-around times. In addition, these NPCs can be used as an *in vitro* platform to study differentiation, cell fate, synaptogenesis and synaptic maturation, as well as electrophysiological properties in a highly controlled environment. These approaches together can provide insight into age-related changes in neural progenitors and facilitate studying cell-autonomous properties that guide adult-born neuron development and integration.

### Comparison with other methods

Most existing protocols were originally developed for, and predominantly applied to, the growth of embryonic and adult SVZ-derived stem cells. However, hippocampal NPCs produce neurospheres that are fewer in number, smaller in size on average, and are more slowly growing than ones isolated from the SVZ^23^. Thus, the methodology traditionally used to isolate and culture neurospheres from the SVZ is not optimal for hippocampal neurosphere growth. The protocol herein was developed to allow for the isolation of NPCs from the DG SGZ of aged (>12 months-old) mice and can be applied to younger animals.

When developing the current protocol, we used the Neural Tissue Dissociation Kit for Postnatal Neuron isolation and found that the use of this kit allows for high cell yield and viability compared to other commonly utilized enzymatic methods (e.g., Papain, Trypsin^24–26^). An important key step we introduced is to initially grow the cells as adherent and let them attach to a coated culture plate. This enables the relatively rare progenitors to proliferate and reach confluence before harvesting and propagation as free floating neurosphere cultures. This protocol offers advantages over methods previously established for isolating cells from the early postnatal and young adult animal tissues, in particular for those researchers interested in investigating the hippocampal neural precursor and progenitor cells in aging.

### Experimental design

The general outline for isolating and culturing NPCs from adult and aged brain includes enzymatic digestion of dissected DG tissue, partial purification from postmitotic neuronal and non-neuronal cells, followed by plating, first as adherent cells on a substrate coated plate followed by culturing as non-attached neurospheres. For optimal protocol outcome the DG dissection and plating should be performed within 2 hours from animal euthanasia. We recommend rapid and gentle handling of the tissue at all steps to ensure maximum cell viability. Collected brain tissue should be kept in the Dissection solution on ice until the dissociation protocol is initiated. Once this protocol is mastered it is possible to perform dissection of 3 to 4 animals at a time.

Before initiating the protocol, researchers should consider whether established neurosphere cultures will be immediately used for downstream applications or will be passaged a fixed number of times before use. While having been shown to increase neural progenitor numbers in culture, passaging may decrease cell viability^24^. This protocol also describes a procedure for neurosphere maintenance in culture. This step should be performed regularly to avoid attachment and differentiation of spheroids. Passaging of neurosphere cultures also helps to avoid excessive cell aggregation due to intrinsic mobility of spheres. It is advisable to freeze multiple vials of established neurospheres to ensure a supply of well-characterized cells. We recommend freezing at least 3 vials per cell line per passage.

Depending on downstream applications, researchers may be interested in changes in neural precursor cultures in a specific disease or condition. For such comparative studies, establishing cultures from individual animals is useful for example, from individual transgenic or wild-type. It is critical to include samples from appropriate controls such as confirming animal genotype before culturing cells and using wild-type and mutant animals from the same litter.

While the current protocol was designed and validated for generating neurospheres from aged mice, we also tested the protocol on early postnatal and young adult animals and were able to successfully establish neurospheres from these time points as well. Therefore, we consider that the current protocol can be utilized for isolating NPCs from a wide range of ages.

### Expertise needed to implement the protocol

A qualified researcher experienced in molecular and cell biology with institutional approval for animal use studies can carry out the complete protocol successfully. A basic understanding of the structure and anatomy of the rodent brain is required to ensure that the correct tissues are collected.

### Limitations

A potential limitation of our protocol is low yield and long culturing times for neurospheres isolated from the aged animals. However, this could be attributed to decrease in the number of NPCs and increased quiescent state of these cells.^27–29^ Long-term cell cultures like the one described in this protocol are prone to mycotic and bacterial infections, which can be remedied by good aseptic technique and the addition of antibiotics/antimycotic to the medium.

Another important consideration is that the NSA might not be effective in detecting and isolating quiescent stem cells. This is because quiescent stem cells are in the G_0_ phase of the cell cycle and therefore not actively dividing. By contrast, this NSA primarily selects for cells that are undergoing proliferation and allows for their expansion *in vitro*.

## Acknowledgements

This work was supported by the National Institutes of Health (NIH) National Institute on Aging grant R01 AG054649 and NIH National Institute of Neurological Disorders and Stroke R21NS115092 to H.-J. Cheng and K. Murray; Academia Sinica grant: AS-GCP-113-L02 to H.-J. Cheng. And NIH National Institute of Mental Health R01 MH119347 (parent grant) and R01 MH119347-S1 (supplement to promote diversity) to E. Diaz.

## Materials

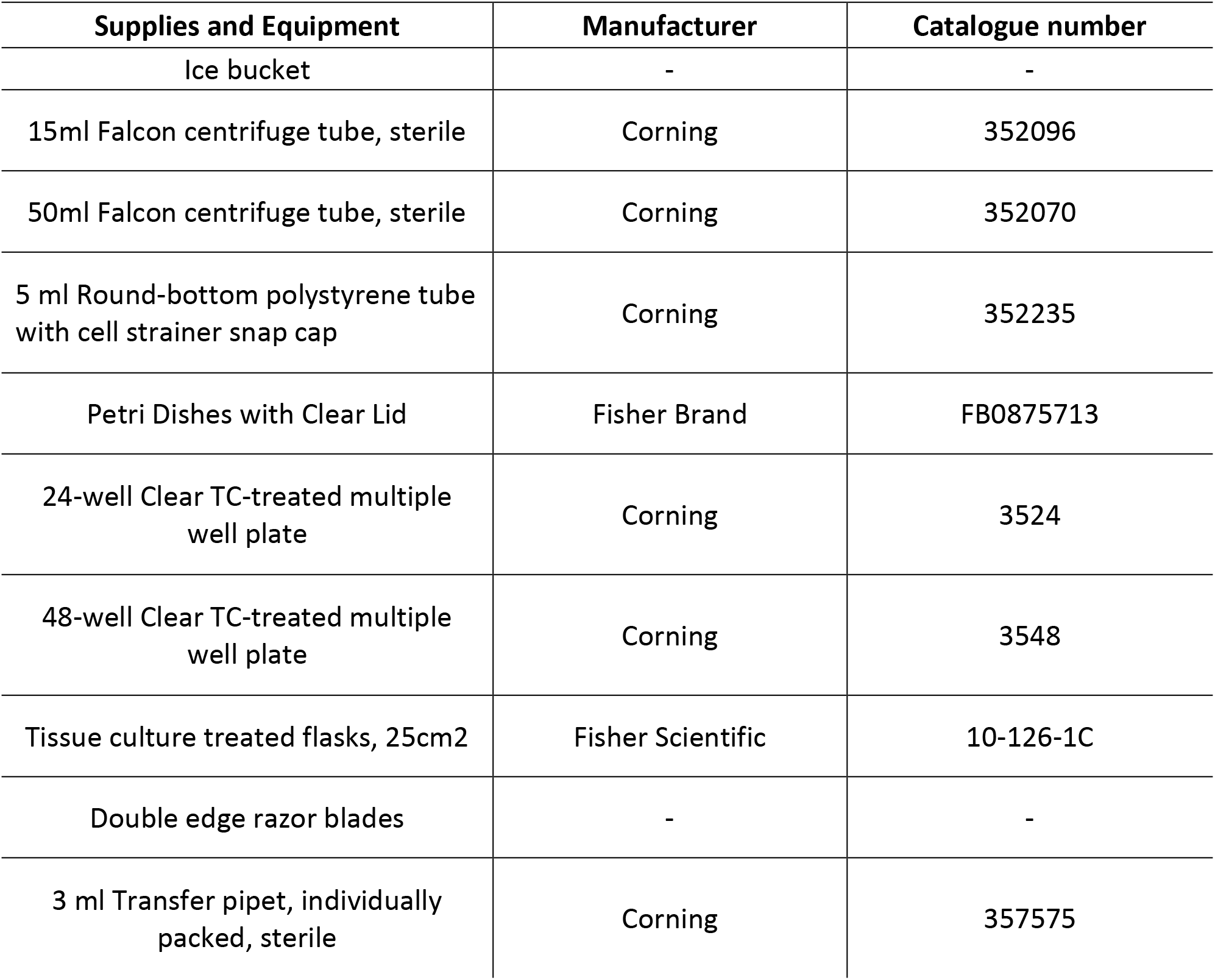
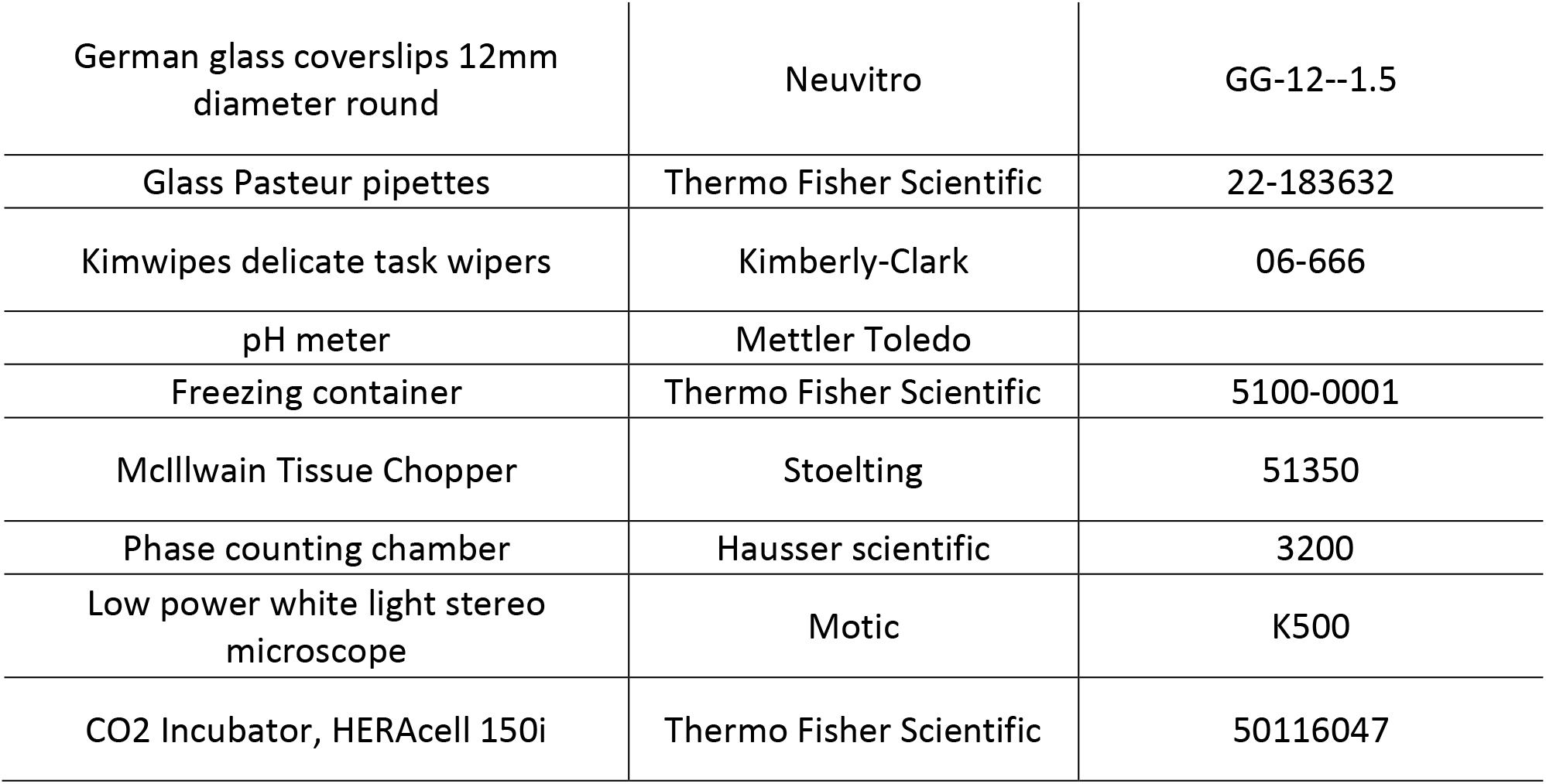

### Animals

| Strain common name | Strain # | Vendor |
| --- | --- | --- |
| C57BL/6J | 000664 | The Jackson Laboratory |
| Gli1-CreERT2 | 007913 | The Jackson Laboratory |
| Ai9 | 007909 | The Jackson Laboratory |

### Dissection tools

| Item # | # |
| --- | --- |
| Light Operating Scissors 5" Straight Sharp/Sharp | 1 |
| Dissector Scissors, Angled to side Sharp /Sharp | 1 |
| Micro Spoon Spatula 5 1/2" | 3 |
| Micro Dissector spatula 5 1/2" | 1 |
| Fine Forceps, Dumont #5 | 2 |

### Reagents and media

| **Reagents** | **Manufacturer** | **Catalogue number** |
| --- | --- | --- |
| Neural tissue dissociation kit<br>(enzyme P) | Miltenyi Biotec | 130-092-628 |
| HBSS, calcium, magnesium,<br>no phenol red | Gibco | 14025092 |
| Sodium bicarbonate<br>(NaHCO<sub>3</sub>) | Sigma-Aldrich | S5761 |
| D-Glucose (Dextrose) | Sigma-Aldrich | G7021 |
| HEPES | Gibco | 15630080 |
| DPBS | Gibco | 14080055 |
| DMEM/F12 | Gibco | 11320033 |
| N-2 Supplement (100X) | Gibco | 17502048 |
| Bovine Serum Albumin | Sigma-Aldrich | A9418 |
| PBS | Gibco | 10010023 |
| Neurobasal medium | Gibco | 21103049 |
| Antibiotic-Antimycotic (100X) | Gibco | 15240096 |
| GlutaMAX Supplement | Gibco | 35050061 |
| B-27 Supplement (50X),<br>minus vitamin A | Gibco | 12587010 |
| B-27™ Supplement (50X),<br>serum free | Gibco | 17504044 |
| Recombinant Human FGF-basic (154 a.a.) | PeproTech | 100-18B |
| Animal-Free Recombinant<br>Human EGF | PeproTech | AF-100-15 |
| (Z)-4-hydroxy Tamoxifen | Cayman Chemicals | 14854 |
| Ethyl alcohol, Pure | Sigma-Aldrich | E7023 |
| Poly-L-ornithine | Sigma-Aldrich | P3655 |
| Laminin Mouse Protein | Gibco | 23017015 |
| Accutase | Stemcell Technologies | 07920 |
| Triton X-100 | Sigma-Aldrich | T8787 |
| TWEEN 20 | Sigma-Aldrich | P1379 |
| Freezing medium Cryo-SFM | Sigma-Aldrich | C- 29912 |
| 2 mL Cryogenic vial | Corning | 431386 |
| Paraformaldehyde solution,<br>8% | Electron Microscopy Sciences | 1578100 |
| Trypan blue solution, 0.4% | Gibco | 15250061 |
| DAPI | Invitrogen | D1306 |
| ProLong Gold Antifade<br>Mountant | Invitrogen | P36930 |
| Microscope Slides | Thermo Fisher | PO103 |
| Collagen, Type 1 rat tail | Corning | 354236 |
| Horse Serum | Gemini Bioproducts | 100-508 |
| Sodium Pyruvate | Invitrogen | 11360-070 |
| MEM | Invitrogen | 51200-038 |
| Papain | Worthington | LS003126 |
| Poly-D-lysine | Fisher | P7405 |
| Trypsin-EDTA | Invitrogen | 25300-054 |
| Sodium tetraborate decahydrate | Sigma-Aldrich | S9640 |
| Boric acid | Sigma-Aldrich | B0394 |

### Antibodies

| Target | Host | Isotype | Clone | RRID | Source | Catalog # | Dilution |
| --- | --- | --- | --- | --- | --- | --- | --- |
| Nestin | Mouse | IgG1 | 10c2 | AB_627994 | Santa Cruz | sc-23927 | 1:500 |
| Sox2 | Rabbit | IgG | EPR3131 | AB_10585428 | ABCAM | ab92494 | 1:10000 |
| GFAP | Mouse | IgG1 | N206A/8 | AB_2877343 | NeuroMab | 75-240 | 1: 1000 |
| MAP2 | Rabbit | IgG | - | AB_91939 | Millipore | AB5622 | 1:750 |

### Tissue dissociation kit

Prepare the Neural tissue dissociation kit (enzyme P) enzyme mix solution as recommended by the manufacturer.

### Dissection solution

1x HBSS

26 mM NaHCO3

30 mM D-Glucose (Dextrose)

2 mM HEPES

Filter-sterilize (0.22 µm) and keep at 4°C. For long-term storage keep in -20°C.

### Dissection medium

495 ml DMEM/F12

5 ml Anti-Anti (100X stock)

Filter-sterilize (0.22 µm) and keep at 4°C.

Aliquot 49.5 ml of dissection medium and add 500 µl of N2 supplement (100X). Keep the medium at 4°C up to 2 weeks.

### Preparation of growth factors

bFGF

EGF

0.1% BSA solution

Reconstitute bFGF and EGF with 0.1% BSA solution in PBS at a concentration of 100 μg/mL. Freeze 20 µl aliquots at -80°C for long term storage. Avoid freeze thaw cycles, keep defrosted aliquot at 4°C up to a week.

### Medium for NPCs maintenance

490 ml Neurobasal medium

5 ml Anti-Anti (100X stock)

5 ml GlutaMAX (100X stock)

Filter-sterilize (0.22 µm) and keep at 4°C up to a month.

Make 49ml aliquots of NPCs maintenance medium. Keep the medium at 4°C up to 2 weeks.

Before use add 1000 µl of B27 supplement-vitamin A (50X) and 10 µl of each bFGF and EGF growth factors on the day of the culture preparation.

Note: bFGF final concentration is 20 ng/ml; EGF final concentration is 20 ng/ml.

### Medium for NPCs differentiation

490 ml Neurobasal medium

5 ml Anti-Anti (100X stock)

5 ml GlutaMAX (100X stock)

Filter-sterilize (0.22 µm) and keep at 4°C up to a month.

Make 49ml aliquots of NPCs maintenance medium. Keep the medium at 4°C up to 2 weeks.

Before use add 1000 µl of B27 supplement (50X) on the day of differentiation experiment.

### Coverslip coating with Poly-L-Ornithine/Laminin

Place 12mm coverslips into 24-well plate. Center the coverslips in the wells; they shouldn’t touch the sides of the well, otherwise the coating solution might spread out outside the coverslip. Pipette 100 µl of 20 ug/ml poly-L-ornithine in DPBS onto each coverslip. Leave the solution overnight. The next day aspirate poly-L-ornithine solution and air-dry coverslips by keeping plate lid off in the biosafety cabinet. The 24-well plates with polyornithine-coated coverslips can be sealed with parafilm and kept at 4°C for a month for future use. A day before use for culture, pour 100 µl of 5 µg/ml laminin in PBS solution per coverslip. Keep the laminin solution on until ready for cell plating, aspirate right before plating cells. Avoid drying out of laminin before cell plating.

### Preparation of 4-hydroxytamoxifen (4-OH TM)

5mg 4-OH TM

250 µl Ethanol, 200 proof.

Dissolve 5 mg 4-OH TM powder in 250 µL molecular grade ethanol to make 1µM stock solution. Gently swirl until completely dissolved. Store at -20°C protected from light.

### Medium for Astrocyte plating and maintaining

434 ml MEM

5ml GlutaMAX

6 ml 30% D-Glucose solution in MEM

5 ml Antibiotic Antimycotic

50 ml Normal horse serum

Prepare 30% w/v % D-Glucose solution in MEM first.

Filter-sterilize (0.22 µm) and keep at 4°C up to a month.

### Borate buffer

Prepare 0.1 M Borate Buffer by dissolving 17.2 g sodium tetra-borate and 3.1 g boric acid in 900 mL of sterile water. Adjust pH to 8.5 and bring the volume up to 1 L with sterile water. Filter-sterilize the solution.

### Poly-D-Lysine stock preparation

To prepare 10x stock of Poly-D-Lysine (10mg/ml) dissolve 100mg bottle in 10ml of borate buffer. Filter-sterilize the stock solution before aliquoting. Aliquot in 1ml aliquots and store in -20C freezer. At the time of coating prepare 1mg/ml poly-D-lysine in borate buffer by diluting 1ml 10X aliquot in 9ml borate buffer.

### Culture flask coating with Poly-D-Lysine

*For coating one 25cm^2^ flask

4ml ddH2O (autoclaved)

400μl collagen

80μl poly-D-lysine (1mg/ml)

Mix all regents together in a 15ml tube. Pour 4480 μl of the coating solution per flask, tilt the flask side to side to evenly distribute the solution. Incubate the flask for 10 min in the biosafety hood, then aspirate the coating solution. Put the lid slight askew and let the flask to dry ∼10-15min. Can be scaled linearly for additional plates.

### Coverslip coating with Poly-D-Lysine

1mL poly-D-lysine (10mg/ml)

9mL Borate Buffer

Prepare 1mg/ml Poly-D-Lysine solution by diluting 10x stock in borate buffer in 15 ml tube. Place 18mm coverslips into 12-well plate. Pipette 250µl of solution per 18 mm coverslip by touching the pipette tip lightly to the surface of the coverslip and ejecting the coating so it forms a bubble on the coverslip. Let the coating solution incubate overnight in the incubator. Next day aspirate coating solution and wash coverslips 2 times with sterile water in the biosafety hood. Leave with water in the wells before ready for cell plating.

*Don’t leave Poly-D-Lysine solution on coverslips for longer than 24 hours-the solution can dry out and form crystals.

### Neural plating media for primary hippocampal culture

432.5 mL MEM

50 mL Donor horse serum

7.5 mL 30% D-glucose stock in MEM

5 mL Sodium pyruvate

5 mL Antibiotic Antimycotic

Filter-sterilize (0.22 µm) and keep at 4°C up to 2 weeks.

## Procedure

### Dissociation and plating of dentate gyrus cells of adult mouse

This step describes the procedure for brain dissection followed by digestion and trituration of tissue into single cells (Fig. 1 A).

**Figure 1.**
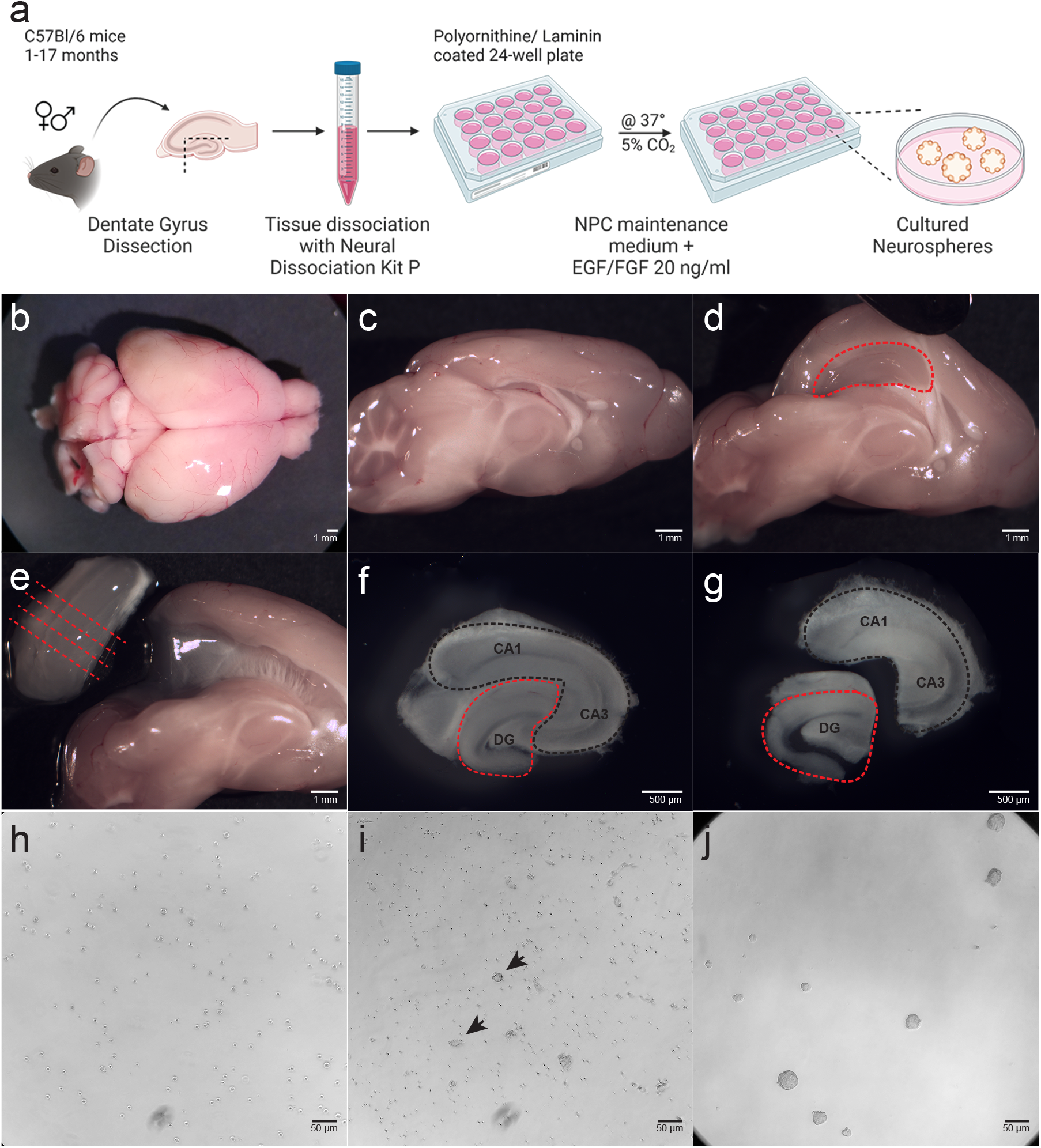
Neurospheres isolated from the dentate gyrus of adult or aged mouse brain. a) Workflow for isolation and culturing of neurospheres from adult or aged mouse dentate gyrus. b) Brain is removed from the skull and placed in dissection dish. c) Hemispheres are separated using a razor blade at the sagittal plane. d) Spatula is used to move the cortex from the midline to free the hippocampus. e) Isolated hippocampus before sectioning in tissue chopper. Red dashed lines indicate the direction of sectioning. f) A slice of hippocampus after tissue chopper. Red dashed lines show outline of dentate gyrus (DG), black lines, cornus ammonis (CA) regions CA1 and CA3. g) Microdissected DG separated from CA regions. h) Bright field image of dissociated DG cells plated after dissection. i) Formation of clusters of newly-formed neurospheres (arrows). j) Neurospheres formed after the first passage. Scale bar: 50 µm.

Timing: ∼2 hours

1. Euthanize an adult mouse by intraperitoneal injection of pentobarbital sodium solution (e.g.,Fatal-Plus solution NDC Code(s): 0298-9373-68) to the dose of >100 mg/kg according to an IACUC -approved protocol. Three-5 min after injection assess euthanasia by tail, toe and eye blink reflex. In the absence of a reflex, decapitate the animal with a large scissors placed just above cervical region of spinal cord.

2. Make a caudal-rostral cut into the skin with the same scissors from the base of the skull toward the nose to open the scalp. Make a straight cut into the skull bone in between eye sockets and two short horizontal cuts on both sides of the skull base.

3. Using small-angled scissors cut the skull in a caudal-rostral direction toward the nose to open skull. Carefully slide a blunt spatula into the cut, moving it underneath the skull bone in between the bone and the brain and push it to the side to open the skull like a “book”. Repeat with other side.

4. Place the spatula underneath the brain moving from the front to back and flipping the brain out of the skull.

5. Put the whole brain in a 50 ml tube containing ice-cold DPBS buffer to wash out the blood.

6. Transfer the entire brain to a 100 mm dish containing ice-cold Dissection solution (Fig.1 B) and make a sagittal cut with a razor blade through the central fissure to separate left and right hemispheres

7. Place a hemisphere lateral side down (Fig.1 C). Place one of the spatulas in the fold between the cortex and the midbrain. Carefully push the cortex to the side exposing subcortical structures. Flatten the cortex with a spatula while using another spatula to hold it in place, hippocampus should be visible in the caudal part on the ventral surface of the cortex (elongated, C-shaped structure) (Fig.1 D). Use one of the spatulas to peal the hippocampus from underneath the cortex. Isolate hippocampi from both hemispheres (Fig.1 E).

8. Hippocampi are sectioned coronally to 500 µm width slices using tissue chopper.

9. Transfer slices into 60 mm dish with ice-cold DPBS. Under a dissection microscope identify the dentate gyrus (DG) and CA regions (Fig.1 F). Use a blunt end spatula to hold a slice and a mini spatula to isolate the DG by cutting between CA1/CA2 and upper blade of DG and making another cut between CA3 and the end of the hilus (Fig.1 G). Perform this procedure on all the slices. Collect DGs in a 15 ml tube containing iced-cold Dissection solution.

10. When tissue slices sink to the bottom of the tube, carefully aspirate Dissection solution. Then add 1950 µl of pre-warmed (37°C) enzyme mix I (1900 µl Buffer Y+50 µl Enzyme P from Miltenyi dissociation kit) and incubate in 37°C water bath for 15 minutes. Invert the tubes about 15 times every 5 minutes.

11. Add 30 µl cold enzyme mix II (20 µl Buffer Y+ 10 µl Enzyme A from Miltenyi dissociation kit) to the sample suspension. Use a fire-polished glass pipette (∼0.8 mm diameter) to gently triturate 10 times. Avoid making air bubbles. Return tubes to the water bath to incubate for additional 10 minutes. Gently invert the sample about 10 times every 5 minutes.

*While waiting for incubation, turn on the centrifuge and set it to 500g; prepare strainer tubes (Corning 35 nm) by pouring 2 ml of prewarmed (37°C) Dissection media through the strainer into the tubes; place NPCs maintenance media into 37°C water bath to prewarm.

13. After 10 minutes of incubation, triturate the cell suspension further using a smaller-diameter (∼0.5 mm) fire-polished glass pipette 10 times.

14. Pipette the cell suspension through the cell strainer into the tubes with Dissection media by placing the tip of the pipette perpendicularly against the strainer mesh and slowly expel the cell solution.

15. Centrifuge the tubes at room temperature at 500g rpm for 5 minutes.

16. Carefully aspirate the media without disturbing the cell pellet, and gently resuspend cells in 3 ml of prewarmed Dissection media and centrifuge the cell suspension again for 5 minutes at 500g.

17. Aspirate the Dissection media and resuspend cells in 500 µl of prewarmed NPCs maintenance medium

18. Remove laminin solution from the coverslips in the prepared plate.

19. Plate the cell suspension into the wells of the plate: 2 hippocampi from 1 brain into 1 well in 24-well plate or 1 hippocampus per well in 48 well plate (2 wells per brain) and put the plate into the incubator.

20. After 2 hours of plating, remove all the media and replace with fresh prewarmed NPCs maintenance medium. Check for cell attachment under the light microscope.

21. Change one third to one half of the NPCs maintenance media every other day.

### Cell culture maintenance procedure

This step describes the procedure for neurosphere maintenance and passaging. It is recommended to perform passaging once every 7 days to keep neurospheres size around 50-100 mm. While all neurospheres are composed of mixed population of precursor cells at various differentiation stages, smaller size help to keep more homogeneous cell composition. Diversity of stem, proliferating neural progenitor cells and postmitotic neurons and glia within spheroids increases with size since more differentiated cell types arise after longer time in culture. Also, cells in the center within bigger spheroids (>250um) suffer from poor gas exchange and receive less nutrients from the cell media. That might lead to cell death ^24^. Cell passaging can be repeated weekly (Fig. 2 A-C), resulting in an exponential increase in total cell numbers. Cells of certain passage number can be cryopreserved for future use as discussed in the next section.

**Figure 2.**
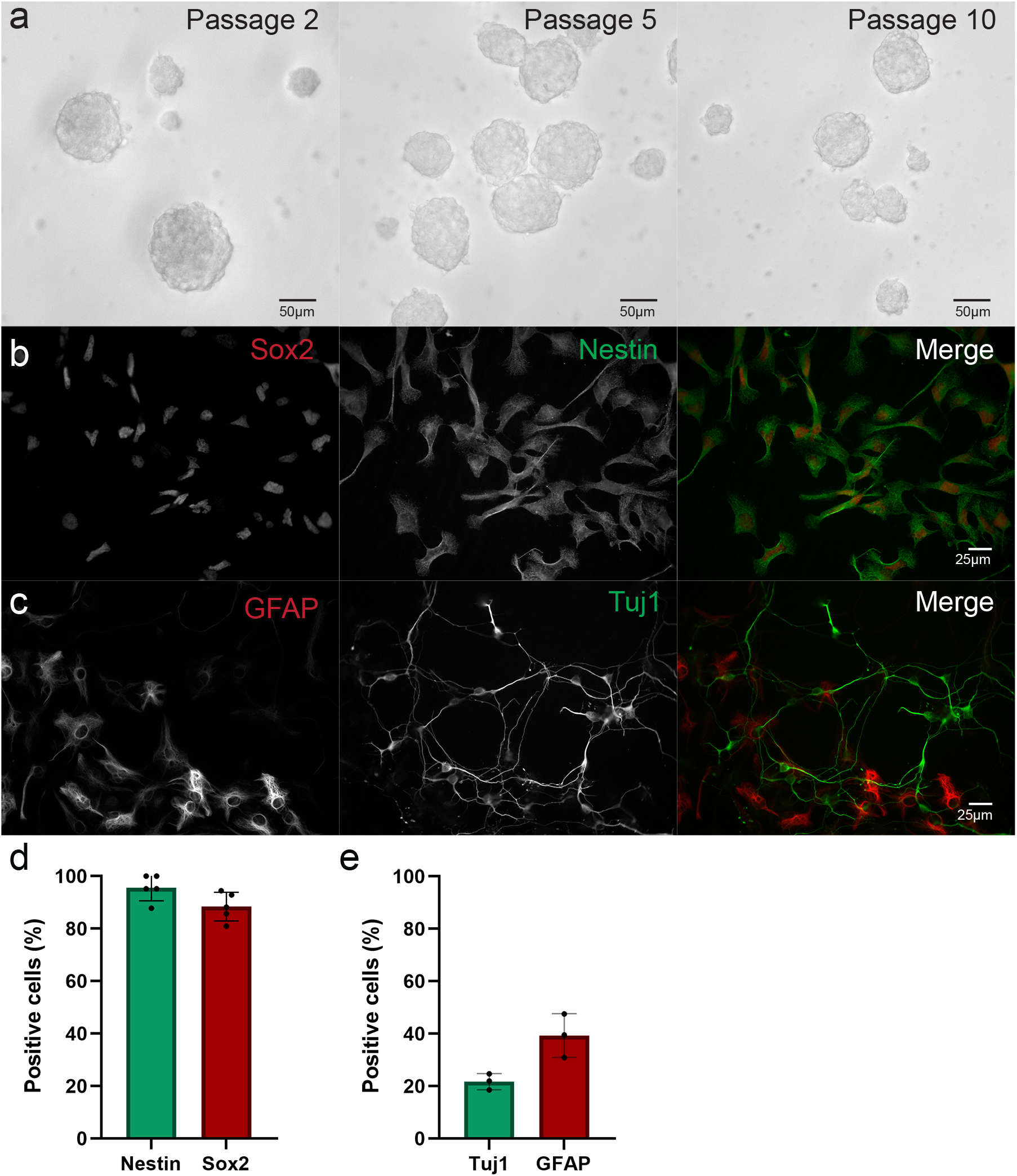
Adult-derived neurospheres exhibit neural progenitor properties. a) Adult-derived neurospheres exhibit capacity for self-renewal. Neurospheres from 11.5-month-old mouse continue to form after passaging cells 2, 5 or 10 times. All images are taken at DIV3 after platting. Scale bar: 50 µm. b) Dissociated NPCs from neurospheres derived from 11.5-months-old mouse at DIV1. The cells were immuno-stained with neural stem cell markers Nestin and Sox2. Scale bar:25 µm. c) Dissociated NPCs from neurospheres derived from 11.5-months-old mouse after 8-day differentiation in culture (DIV8) differentiated. The cells were immuno-stained with astrocytic marker GFAP and for neuronal marker Tuj1. Scale bar: 25 µm. d) Quantification of Nestin and Sox2 expressing cells at DIV1 post plating. e) Quantification of GFAP and Tuj1 expressing cells at DIV8 post plating.

22. Prewarm NPCs maintenance medium in the water bath to 37°C.

23. Use a P1000 pipette to transfer contents of the well to be passaged into a 15 ml tube. If there are neurospheres attached to the bottom of the well, add fresh 500 µl of NPCs maintenance medium to the well and use the p1000 pipette to gently dislodge these cells by pipetting up and down. Collect the medium and combine with the rest of well contents.

24. Spin the tube at 500xg for 5 min at room temperature.

25. Aspirate medium without disturbing the pellet.

26. Add 500 µl of prewarmed Accutase enzyme mix to the tube and use p1000 pipette to resuspend the pellet by triturating 3-4 times.

27. Place neurosphere suspension at 37°C water bath for 5 minutes.

28. Spin down cells at 500xg for 5 minutes at room temperature.

29. Remove supernatant.

30. Resuspend the cells in 1 mL of NPC maintenance medium + growth factors and calculate the cell concentration using counter chamber.

31. Plate cells in 500 µl of NPCs maintenance medium with growth factors at the concentration of 1x10^5^ - 5x10^5^ cells/mL in an uncoated 24-well plate.

32. Small neurospheres should be observed by the next day.

33. Change 1/3 of medium every other day.

It is important to monitor cultures every day to determine the conditions of the neurospheres. The neurospheres should appear as round with light edges with small protruding cilia and translucent/not dark center. Cell culture medium should be pink colored. In case if the medium appears yellow the frequency of media change should be adjusted (e.g., be everyday) to keep cell healthy.

### Preparing a frozen stock of NPCs

At passaging, neurospheres can be frozen and stored in liquid nitrogen for future use, which is described here. Please note that steps #34-36 are identical to steps #22-24 in the previous section.

34. Use a p1000 pipette to transfer contents of the well to be passaged into a 15 ml tube. If some neurospheres attached to the bottom of the well, add 500 µl of fresh NPCs maintenance medium to the well and use the p1000 pipette to gently dislodge these cells by pipetting up and down. Collect the medium and combine with the rest of the well contents.

35. Spin the tube at 500xg for 5 minutes at room temperature.

36. Aspirate medium without disturbing the pellet.

37. Add 1000 µl of room temperature Cryo-SFM medium to the tube and use a p1000 pipette to triturate and resuspend the pellet. Avoid introducing air bubbles.

38. Transfer the cell suspension to a cryovial previously labeled with information about the neurosphere cell line, passage number and the date of freezing.

39. Place the cryovial into a freezing container.

40. Put the container with cells at -80°C freezer for at least 8 hours before transferring vial to the liquid nitrogen cryostorage unit.

### Recovering neurospheres from a frozen stock

This section describes procedure for recovery of cryopreserved neurosphere cell lines.

41. 2 days prior to the planned recovery date start to prepare 24-well culture plate by wells with 300 µl of 20 ug/ml poly-L-ornithine in DPBS into each well. Leave the solution overnight.

42. The next day aspirate poly-L-ornithine solution and air-dry in the coverslips by keeping plate lid off in the biosafety cabinet.

43. Add 300 µl of 5 µg/ml laminin in PBS solution per well. Keep the laminin solution in a well until ready for cell plating, aspirate right before plating cells. Avoid drying out of laminin before cell plating.

44. Retrieve cryovials with frozen neurospheres and promptly place into a water bath at 37°C.

45. Keep the cap of a cryovial above water when swirling the cryovial to prevent contamination from the water bath. Swirl the vial to thaw contents evenly.

46. When the cell solution is fully thawed, remove the cryovial from the water and spray thoroughly with 70% ethanol before placing it into the biological safety hood.

47. Slowly in dropwise fashion add 1000 µl of pre-warmed NPCs maintenance medium per cryovial. Carefully pipette up and down 1-2 times

48. Transfer cell solution to a 15 ml tube and spin it down at 500g for 5 minutes at room temperature.

49. Aspirate medium and resuspend cell pellet in 500 µl prewarmed NPCs maintenance medium per tube.

50. Remove laminin from the well and add neurosphere suspension into culture plate.

51. 4-6 hours after plating check the plated cells under the light microscope. Some neurospheres should attach to the bottom of the plate and cells migrating out of attached spheroids should be visible.

52. Change 200 µl of the medium every other day.

### Plating neurosphere cells for differentiation

In this step, we describe differentiation of the neuroshere cells plated on the coverslips.

Timing: 4-5 days

53. Two days before plating cells for differentiation, prepare Poly-L-Ornithine/Laminin-coated coverslips as described in the Materials section.

54. On the day of the experiment, prewarm a 50ml aliquot of NPCs maintenance medium to 37°C.

55. Check the culture. Neurospheres should appear round with light bright edges with small protruding cilia and translucent/not dark center under the light microscope.

56. Collect media with neurosheres from a well of a 24-well plate into a 15 ml tube.

57. Spin the tube at 500g for 5 minutes at room temperature.

58. Aspirate supernatant without disturbing the cell pellet.

59. Add 500 μl of Accutase prewarmed to room temperature.

60. Incubate at for 5 minutes in the biosafety hood at room temperature.

61. Add 500μl of 37°C NPCs maintenance medium and collect with a p1000 pipette.

62. Spin the tube at 500g for 5 minutes at room temperature.

63. Remove supernatant and resuspend neurospheres in 1000μl of NPCs maintenance medium.

64. Use fire polished glass pipette to triturate neurosheres until they are dissociated into a single-cell solution, between 5 to 10 times.

65. Pipette 10μl of dissociated cell solution into a 1.5 ml tube. Add 10μl of Tripan blue solution, mix by pipetting up and down.

66. Load into a hemocytometer and perform live cell count under light microscope.

67. Calculate volume to obtain desired plating density.

### Critical step

Plating density should be optimized for different cell lines, typically, 5x10^4^-1x10^5^ cells per 12 mm coverslip.

68. Plate the desired number of cells onto precoated coverslips with 500μl of NPCs maintenance medium and place in the incubator for 15 minutes for attachment.

69. Check cell attachment at the microscope by gently shaking the plate, attached cells should not float.

70. Change medium to differentiation medium after 2 days *in vitro* (DIV) counting the day of plating as DIV1.

71. Maintain differentiating cells by changing 200 μl of differentiation medium in each well every other day.

### Immunocytochemical staining for NPC, neuron and astrocyte markers

In this step, we describe immunofluorescence-based assays to confirm NPCs identity before using the cells in experimental assays as well the differentiation of these cells to neurons and astrocytes.

### For NPC marker staining

72. Dissociate and plate neurosphere cells as described in steps 53-70

73. After 1 day of differentiation, aspirate the media and rinse coverslips with PBS.

74. Prepare a fresh batch of 4% paraformaldehyde (PFA) by diluting 8% PFA from the ampoule with PBS. Add sucrose to the PFA to make 4% solution by weight. Check pH, adjust to

74. if needed. Prewarm the solution in 37°C water bath for 5 min.

*Alternatively, 4% PFA solution can be prepared in-house. Aliquoted PFA should be stored at - 20°C and are good to use for up to 6 months after preparation. Avoid freeze-thaw cycles.

75. Aspirate the medium and add pre-warmed 4% PFA /4% sucrose solution.

76. Incubate the coverslips in the fixative solution for 12 minutes.

77. Remove the fixative and wash coverslips with PBS 3 times, 5 minutes each.

78. Add permeabilization solution (0.25% Triton X-100 in PBS) for 10 minutes immediately after removing the final wash, avoid drying of the coverslips.

79. Following permeabilization, cells are washed in PBS, once for 30 seconds and then twice for 5 minutes.

80. Incubate the coverslips in blocking solution (5% bovine serum albumin [BSA] in PBS + 0.2% Tween-20 [PBST]) for at least 35 minutes at RT.

81. Apply primary antibodies for neural stem cell markers such as anti-Nestin and anti-Sox2 diluted in blocking solution (5% BSA in PBST) and incubate for 2 hours at room temperature or overnight at 4°C fridge.

82. Remove antibody and wash coverslips with PBS 3 times for 5 minutes.

83. Add appropriate secondary antibodies conjugated to fluorescence probes in the blocking solution and incubate for 1 hour at RT in the dark.

84. Remove the antibody solution and wash coverslips with PBS 3 times for 5 minutes.

85. Stain with DAPI diluted in PBS for 3 min.

86. Wash once with PBS.

87. Mount coverslips with a drop of hard-set mounting medium (e.g., ProLong Gold Antifade Mountant) onto glass slides.

88. Keep the slides in a dark, dry place for at least 2 hours before imaging.

89. Approximately >90% of all cells should be positive for Nestin and Sox2 (Figure 3B and 3C).

**Figure 3.**
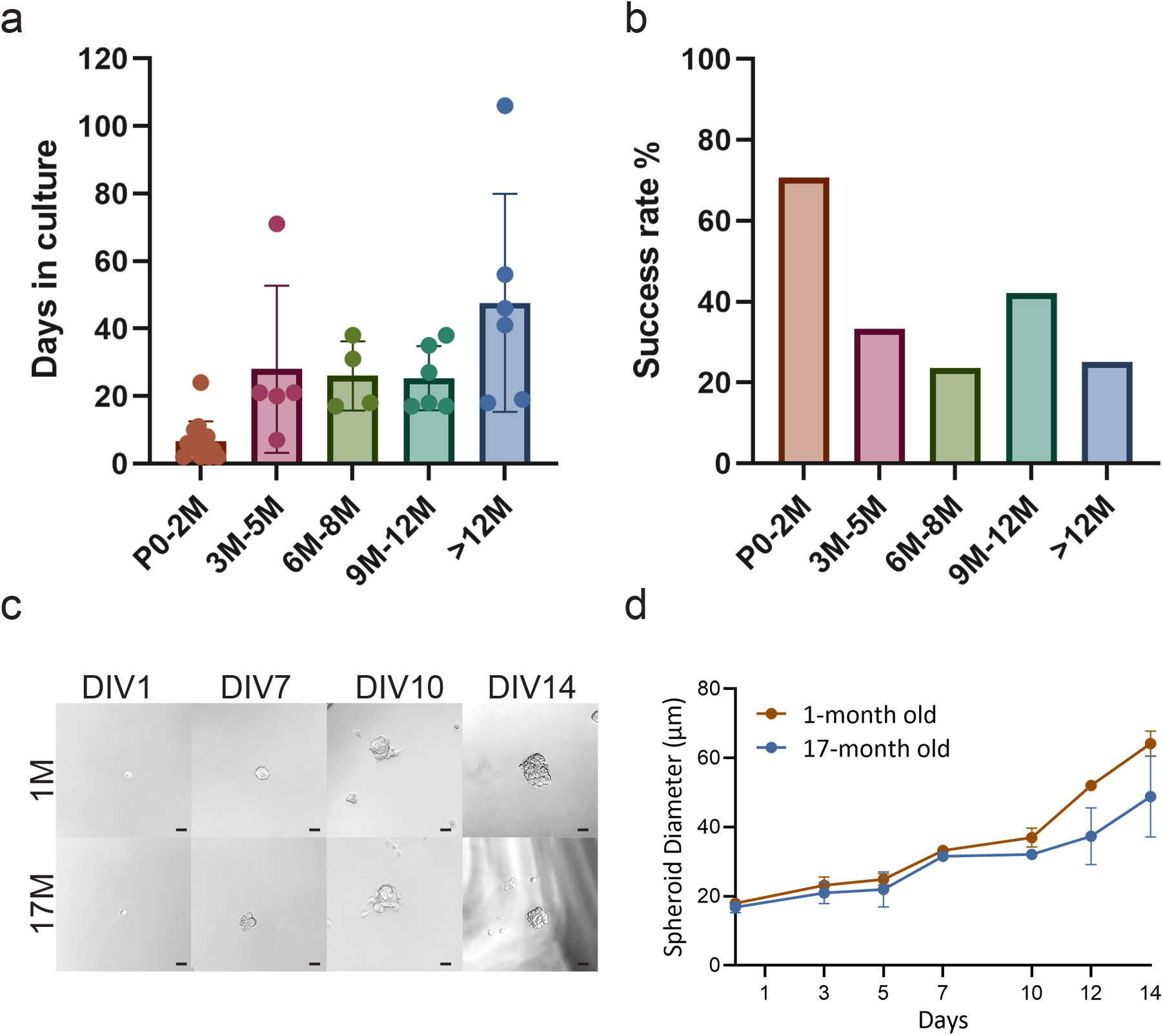
Stable neurospheres can be isolated from adult mice at different ages. a) Time to the formation of neurospheres varies with age of animals, though stable cultures were established at several timepoints between 1 and 12+ months of age. b) Success rate of establishing stable neurosphere cell line decreases with age. Percent calculated by dividing number of neurosphere cell lines established by total number of animals dissected at each indicated time point. c) Brightfield images of growing neurospheres from young and aged animals across 14 days *in vitro* (DIV14). Scale bar: 25 µm. d) Neurospheres from the young and aged animals grow in culture similarly in diameter up to DIV10. However, at DIV12 and 14 neurospheres from aged mice show lower spheroid diameter.

### For staining differentiated neurospheres for neuronal and astrocytic markers

90. Follow steps described in steps 53-70 for neurosphere dissociation and plating.

91. Maintain cells DIV7, by changing 200μl of NPCs differentiating media every over day.

92. After 7 days of differentiation, aspirate the media and rinse coverslips with PBS.

93. Fix and perform immunocytochemical staining as outlined in steps 74 through 88.

94. For step 81, apply primary antibodies such as anti-MAP2 (AB5622, 1:750) and anti-GFAP (NeuroMab N206A/8 1:1000) diluted in the blocking solution (5% BSA in PBST) and incubate for 2 hours at RT or overnight at 4°C.

95. The percentage of MAP2 positive derived neurons and GFAP positive derived astrocytes are each roughly 10%–30% of total cells (Figure 3D).

Differentiated cells can be analyzed by other quantitative methods such as immunofluorescence-based assays, qPCR and flow cytometry.

### *In vitro* tamoxifen-induced recombination for conditional gene expression in neurospheres derived from transgenic mice

This approach can be applied to drive conditional expression of transgene in NPC cell lines by treating neurospheres with 4-OH tamoxifen (TM) (Fig.4 B). Here we show how this protocol can be applied to NPC derived from Gli1CreERT^2^xAi9. In brief, the Gli1CreERT^2^xAi9 mouse line has both (1) a tamoxifen (TM)-inducible form of Cre-recombinase driven by the Gli1-promoter, a gene which is active in neural stem and precursor cells (NPCs), and (2) a loxP site before a stop codon before the Rosa26-tdTomato gene (Fig.4 A). Administration of TM allows for the expression of tdTomato (tdTom) fluorescent protein in cells with the active Gli1 promoter and permanently labels their progeny (Fig.4 C). This allows to identify and track differentiating cell derived from transgenic neurosheres.

**Figure 4.**
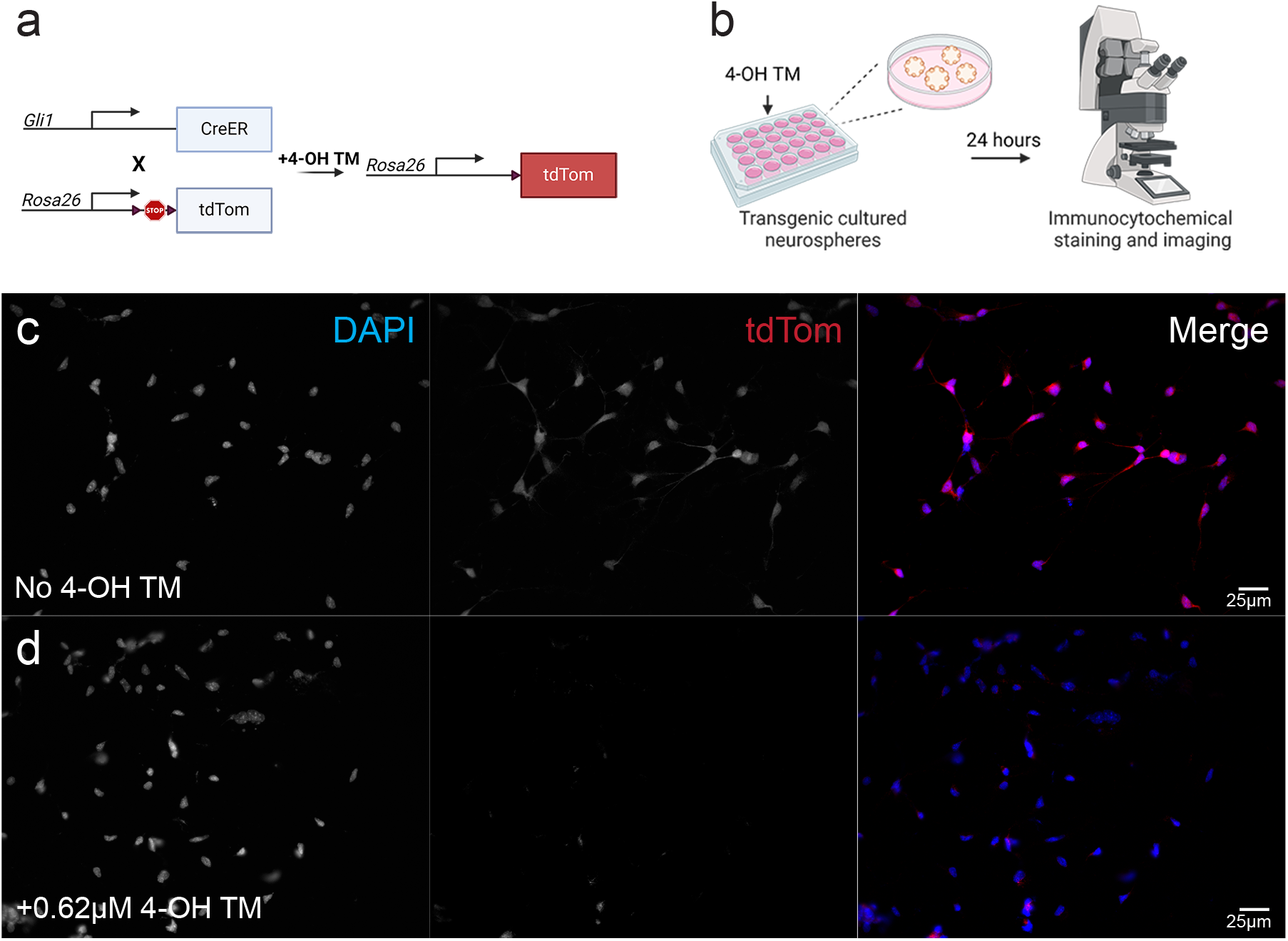
Neurospheres generated from adult or aged transgenic Gli1CreER x Ai9 mouse. a) A schematic diagram depicting how the Gli1CreER x Ai9 mouse was generated. *In vitro* administration of 4-Hydroxytamoxifen (4-OH TM) to the neurospheres derived from these transgenic mice enables tracking of these isolated cells. b) Workflow for *in vitro* induction of neurospheres derived from adult or aged Gli1CreER x Ai9 animals. c) Epifluorescent images of dissociated Gli1Cre x Ai9 neurospheres without 4-OH TM treatment as a control. Cells were immunostained for tdTommato (tdTom) and DAPI. d) Epifluorescent images of dissociated Gli1Cre x Ai9 neurospheres treated with 0.62µM 4-OH TM for 24 hours. Cells were immunostained for tdTom and DAPI. Scale bars: 25µm.

96. Plate 1*10^5^ cells per well cells per 24-well plate following steps 22-33

97. Maintain neurospheres, allow them to grow for 3 days, or until they reach an average diameter of 50-100mm.

98. Prepare media for 4OH-TM induction by dissolving 4OH-TM stock solution in freshly made NPC maintenance media. Final 4-OH TM concentration in the media should be 0.62 µM.

99. Transfer neurospheres from one well of 24-well plate in to 15ml tube and centrifuge it for 5min at room temperature at 500g.

100. Aspirate the supernatant and gently resuspend neurospheres in 500µl NPC maintenance media with 4-OH TM.

101. Return cells into the incubator for 24 hours. After incubation period, NPCs are ready to be used for downstream experiments, e.g., differentiation or co-culture with primary hippocampal neurons in adult-neurogenesis in vitro assay described in the next section.

102. Collect the 4-OH TM induced cells, dissociate and plate them onto glass coverslips as described in steps 53-71.

*We recommend performing 4-OH TM induction before each experiment on freshly expanded neurospheres. That provides a good quality cell labeling highlighting cell morphology. The TdTomato fluorescent protein expression was observed in induced neurospheres and differentiated cells after passaging. However, the labeling appears uneven throughout the cell.

### Heterochronic co-culture of primary hippocampal neurons with neuroshere-derived cells

This section describes how to prepare and maintain early postnatal mouse primary hippocampal culture. Neurospheres can be dissociated and added to the primary culture at DIV7. When added to primary culture, neural precursor cells from neurospheres generate neurons and astrocytes (Fig.5 A). This co-culture method offers a simplified *in vitro* model of adult neurogenesis as neural precursor cells differentiate and mature in the environment of preformed neuronal network of primary culture. We used neurospheres isolated form Gli1CreERT^2^xAi9 mouse line. That allows to identify and track differentiating cell derived from transgenic neurosheres.

**Figure 5.**
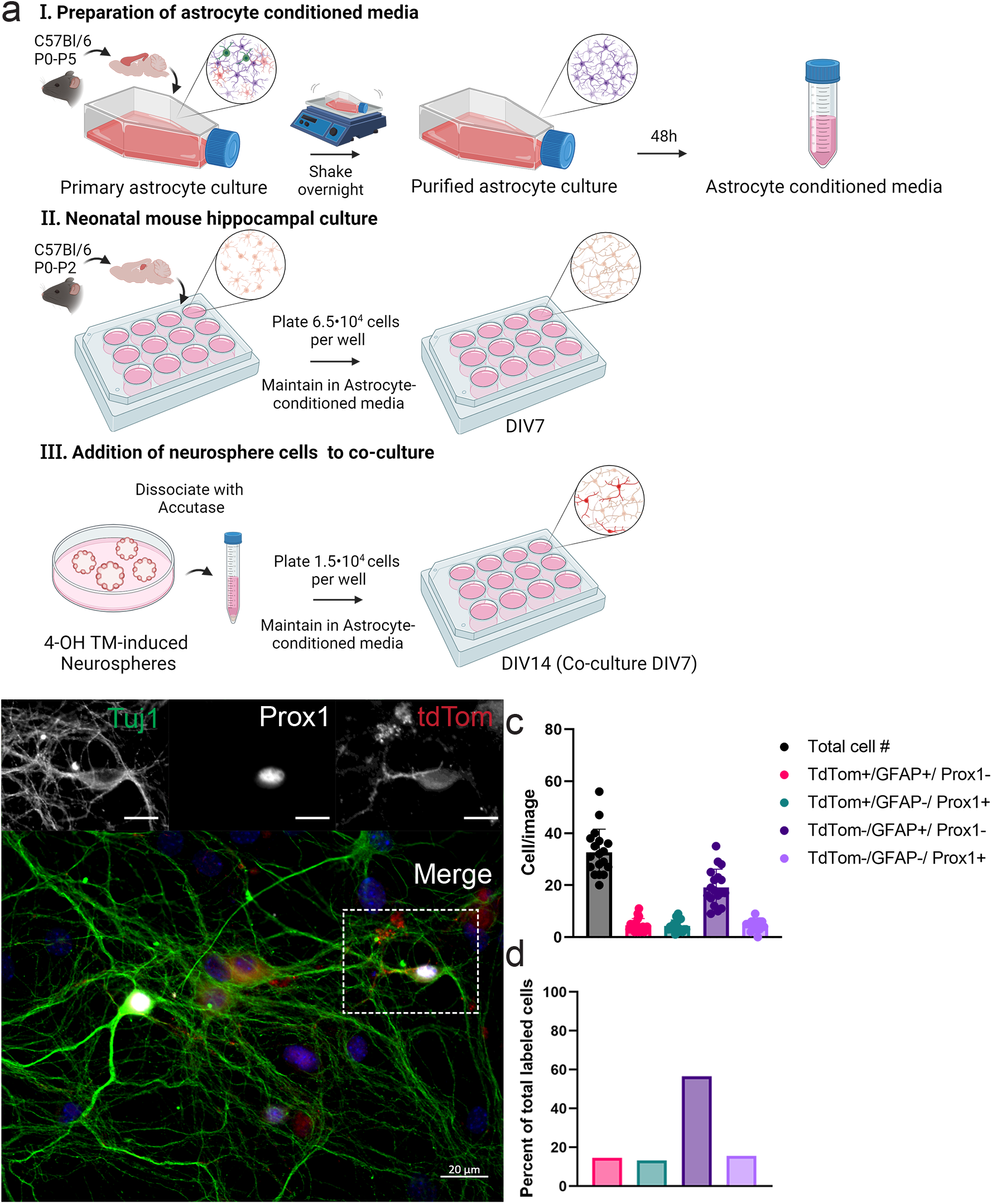
Heterochronic co-culture of granule cells differentiated from neurospheres derived from adult dentate gyrus with hippocampal cells from primary culture. a) Flowchart of preparation of heterochronic co-culture of neurosphere-derived cells with primary hippocampal culture. b) Epifluorescence image of DIV7 co-culture of granule cells and astrocytes differentiated from neurosphere cells derived from 11.5-month-old mouse and primary hippocampal cells. Culture is immunofluorescently labeled for Tuj1, tdTommato (tdTom) and Prox1. Nuclei are counterstained with DAPI. The upper images are from the rectangular area in the bottom image which is a merged image. Scale bar: 20 µm, inset scale bar: 5 µm. c) Co-culture composition shown by combination of tdTom labeling cells derived from neurospheres, astrocytic marker GFAP and marker for granule cells Prox1. d) Percent breakdown of co-culture composition. The combinations of the expression of markers are used to identify specific cell type and origin: tdTom+/GFAP+/ Prox1- : Astrocytes derived from neurosphere culture; tdTom+/GFAP-/ Prox1+: granule cells derived from neurosphere culture; tdTom-/GFAP+/ Prox1-: Astrocytes from primary hippocampal culture; tdTom-/GFAP-/ Prox1-: Granule cells from primary hippocampal culture.

For this protocol we use astrocyte-conditioned medium for maintenance of postnatal mouse primary hippocampal culture as that has been show to promote neuronal health^30^. Prepare astrocyte conditioned medium before postnatal mouse primary hippocampal culture. Conditioned medium can be stored at 4°C for up to month.

### Preparation of astrocyte conditioned media

103. Prepare Poly-D-lysine coated flasks as described under Materials section.

104. Prepare and prewarm Astrocyte plating and maintenance media (15ml media per flask).

105. Mix up a digestion solution by combining 4900µl HBSS + 100µL papain into 15ml tube for each pup to be dissected.

106. Anesthetize the P0-P5 mouse pups by placing them on a kimwipe-covered ice until not responsive.

107. Place new paper towel down and wipe each pup head and neck with ethanol well before rapidly decapitating using sterile scissors.

108. Wrap up body and place in a waste bag. Place a head into a petri dish filled with cold Dissection solution.

109. Cut away skin and peel back to expose skull.

110. Cut into cerebellum and cut along middle of skull: up and out. Make a cut at front at the olfactory bulbs and 2 cuts at the cerebellum, such that the skull can flap open like a book.

111. Peel back skull and scoop out brain with spatula and place in Dissection solution.

112. Look at brain under dissection microscope-use angled forceps to pin brain to bottom of dish through the cerebellum. Use another pair of forceps to remove pia. Start away from the region of interest and go down the midline of one hemisphere “nipping” at pia. Should be able to peel away pia in a sheet.

113. Use microspatula dissector to remove cortex from both hemispheres.

114. Cut cortex tissue into smaller pieces by pinning it down with forceps and cutting with microdissector.

115. Using a sterile plastic pipette to transfer brain tissue to clean 15ml tube. Keep it on ice with a little bit of dissection solution while finishing dissection of other pups.

116. Take tubes with tissue to the biosafety hood. Aspirate dissection solution.

117. Add 5ml of digestion solution per tube.

118. Incubate tubes for 15 minutes at 37°C in water bath. Swirl tubes every 5 minutes.

119. Once the incubation is complete, remove digestion solution.

120. Add 5mL of Astrocyte plating medium to each tube and use a fire-polished Pasteur pipette triturate tissue 5-10 times.

121. Centrifuge the tubes at 500g at room temperature for 5 min.

122. Check for the pellet, suck out the media and resuspend pellet in 1 ml of prewarmed Astrocyte media.

123. Mix 10 µL of cell suspension with 10µL of Trypan blue. Apply 10µL of solution to hemocytometer and count cells.

124. Plate 2.25x10^6^ cells per 25cm^2^ PDL-coated flask in 5mL Astrocyte medium

125. Loosen cap slightly to allow gas exchange in incubator.

126. Change media (5mL) after 3 hours and assess attachment.

127. Change media (5mL) completely next day.

128. Change media every three days until ∼80% confluent.

129. When desired confluency is achieved, shake astrocyte flask(s) at 250 RPM overnight in bacterial shaker in a sealed ziplock bag (flasks should be attached with tape to the bottom of the bag and firmly taped to the table of the shaker.

130. Retrieve flasks from the shaker. Spray down bag with 70% alcohol to avoid contamination. Remove flask from the bag and check under microscope. Astrocytes should be attached to the bottom of the flasks while most of there cells will be floating in the media. Foam in the culture media is normal.

131. Pre-warm 10 ml of Astrocyte media and warm up 1ml Trypsin-EDTA per astrocyte flask to be re-plated.

132. For each flask that will be replated, prepare 3 new 25cm2 flasks by adding 5 mL of Astrocyte medium and place in the incubator to equilibrate.

133. Remove media from the astrocytes and rinse the cells with 5mL of sterile PBS by tilting the flask multiple times.

134. Remove PBS and add 1ml pre-warmed to 37C° Trypsin-EDTA.

135. Incubate for at least 5 mins in the incubator. Check under the microscope for detachment. If cells are still attached, return to the incubator for another minute.

136. Add 9 ml prewarmed Astrocyte media to the flask. Use a 5ml serological pipette to gently wash cells off the bottom of flask. Transfer cell solution to a new 15ml tube.

137. Centrifuge the cells at 500g for 5 minutes.

138. Carefully remove supernatant with vacuum aspirator and add 1ml of Astrocyte media to the pellet and resuspend by triturating the pellet approximately 5x or more if visible aggregates of cells remain in solution.

139. Perform a cell count with 10ul of cell suspension.

140. Plate 3.5x 10^5^ cells per new 25cm flask with 5mls of Astrocyte medium.

141. Change media after 24 hours.

142. Replace media every 3 days until cells are 80-90% confluent.

143. Replace Astrocyte medium with 12.5 ml of NPCs differentiation medium.

144. After 48 H Collect and filter astrocyte conditioned media with a 0.22µm filter (label with date name and batch info: original culture date, split date)

145. Store conditioned media at 4°C for up to one month.

### Primary hippocampal culture

146. Prepare Poly-D-lysine coated coverslips as described under Materials section.

147. Prepare and prewarm Neural plating media to 37C°.

148. Mix up a digestion solution by combining 4900µl HBSS + 100µL papain into 15ml tube for each pup to be dissected.

149. Anesthetize the P0-P2 mouse pups by placing them on a kimwipe-covered ice until not responsive.

150. Place new paper towel down and wipe each pup head and neck with ethanol well before rapidly decapitating using sterile scissors.

151. Carefully remove the brains and place in petri dishes filled with Dissection buffer.

152. Remove the hippocampi from both hemispheres of each brain as described in step 7.

153. Transfer tissue to a 15 mL conical tube on ice with transfer pipette. Keep it on ice with a little bit of dissection solution while finishing dissection of other animals.

154. Take tubes with hippocampal tissue to the biosafety hood. Aspirate dissection solution.

155. Add 5ml of digestion solution per tube.

156. Incubate tubes for 15 minutes at 37°C in water bath inverting gently every few minutes.

157. After incubation in complete, add 5 mL of prewarmed Neural plating media.

158. Spin tubes at 500g at room temperature for 5 minutes.

159. Aspirate the supernatant and add 1 mL of Neural plating media.

160. Dissociate tissue with fire-polished glass pipette by triturating up and down 5-10 times, until no chunks of tissue are visible.

161. Use a hemacytometer to perform live cell count.

162. Retrieve washed Poly-D-lysine coated coverslips form incubator. Aspirate water.

163. Plate cells at 6.5x10^4^ cells in 1.5ml of media per 18mm coverslip in 12-well plate.

164. Place plates in incubator for 4 hours.

165. After 4 hours, completely remove Neural plating media and replace it with Astrocyte conditioned media.

166. Perform media change every 2 days by removing 300µl old media and adding 300µl fresh Astrocyte conditioned media.

### Introducing neurosphere cells to co-culture

167. On DIV7 (7 days i*n vitro*) of primary hippocampal culture collect and dissociate neurospheres as described in steps 55-62 of “Plating neuroshere cells for differentiation” section.

168. If the neurospheres from the inducible Cre system are used in the experiment, the 4-OH TM-induced recombination step described in previous section should be performed before the neurospheres are introduced to the co-culture.

169. After dissociation, resuspend neurospheres in 1000μl of Astrocyte conditioned media.

170. Use fire polished glass pipette to triturate neurosheres until they are dissociated into a single-cell solution, between 5 to 10 times.

171. Perform a live cell count by mixing cell solution with Trypan blue 1:1 and quantifying live cells in hemocytometer.

172. Dilute cell solution to 5x10^5^ cells per ml of Astrocyte conditioned media.

173. Retrieve plates with primary hippocampal culture from the incubator. Remove 300μl of media form each well to be used for co-culture.

174. Slowly, in drop-wise fashion add 300μl dissociated neurosphere cell solution into wells with primary neuronal culture.

175. Return plates into incubator.

176. Perform media change every 2 days by removing 300 old media and adding 300µl fresh Astrocyte conditioned media. Maintain cell culture for another 7 days.

## Troubleshooting

Troubleshooting:
| Problem | Possible reason | Solution |
| --- | --- | --- |
| Bacterial or mycotic contamination | Poor aseptic technique | Include Anti-Anti to the culturing medium. It is advisable to freeze as multiple vials of established neurospheres to ensure a supply of well-characterized cells. Freezing of at least 3 vials per cell line per for each passage number is recommended. |
| Low cell yield after digestion | Enzymatic digestion was insufficient | Increase incubation time while visually inspecting the progress of digestion |
| | Under digestion due to excessive amount of tissue used | Dissociate maximum of 2 hippocampi in 50μl of Enzyme P dissolved in prewarmed 1900μl of Buffer X. |
| Low cell viability after digestion | Excessive trituration | Use fire polished Pasteur pipettes with not sharp edges. Make sure the opening is not too narrow ($\geq 0.5$mm in diameter). |
|  |  | Pipette cells gently and avoid introducing air bubbles |
|  | Harsh handling of cells during cell filtration | Do not scrape the mesh of the strainer when filtering cell suspension to avoid damaging the cells |

## References

1. Jensen JB, Parmar M. Strengths and limitations of the neurosphere culture system - Jensen and Parmar Mol Neurobiol 2006 (1).pdf. 2006;34(3):153-154. file:///C:/Users/marrac/Downloads/Jensen and Parmar Mol Neurobiol 2006 (1).pdf

2. Azari H, Reynolds BA. In vitro models for neurogenesis. Cold Spring Harb Perspect Biol. 2016;8(6):1–11. doi:10.1101/cshperspect.a021279

3. Reynolds BA, Weiss S. Generation of neurons and astrocytes from isolated cells of the adult mammalian central nervous system. Science. 1992;255(5052):1707-1710. doi:10.1126/science.1553558

4. Ming G li, Song H. Adult Neurogenesis in the Mammalian Brain: Significant Answers and Significant Questions. Neuron. 2011;70(4):687–702. doi:10.1016/j.neuron.2011.05.001

5. Altman J, Das GD. Autoradiographic and histological evidence of postnatal hippocampal neurogenesis in rats. J Comp Neurol. 1965;124(3):319–335. doi:10.1002/cne.901240303

6. Palmer T.D., Takahashi J, Gage FH. The adult hippocampus contains primordial stem cells. Mol Cell Neurosci. 1997;8:389–404.

7. Toni N, Laplagne DA, Zhao C, et al. Neurons born in the adult dentate gyrus form functional synapses with target cells. Nat Neurosci. 2008;11(8):901–907. doi:10.1038/nn.2156

8. Saxe MD, Battaglia F, Wang JW, et al. Ablation of hippocampal neurogenesis impairs contextual fear conditioning and synaptic plasticity in the dentate gyrus. Proc Natl Acad Sci U S A. 2006;103(46):17501–17506. doi:10.1073/pnas.0607207103

9. McAvoy KM, Scobie KN, Berger S, et al. Modulating Neuronal Competition Dynamics in the Dentate Gyrus to Rejuvenate Aging Memory Circuits. Neuron. 2016;91(6):1356–1373. doi:10.1016/j.neuron.2016.08.009

10. Hernández-Mercado K, Zepeda A. Morris Water Maze and Contextual Fear Conditioning Tasks to Evaluate Cognitive Functions Associated With Adult Hippocampal Neurogenesis. Front Neurosci. 2022;15(January):1–16. doi:10.3389/fnins.2021.782947

11. Berdugo-Vega G, Arias-Gil G, López-Fernández A, et al. Increasing neurogenesis refines hippocampal activity rejuvenating navigational learning strategies and contextual memory throughout life. Nat Commun. 2020;11(1):1–12. doi:10.1038/s41467-019-14026-z

12. Kuhn HG, Dickinson-Anson H, Gage FH. Neurogenesis in the dentate gyrus of the adult rat: Age-related decrease of neuronal progenitor proliferation. J Neurosci. 1996;16(6):2027–2033. doi:10.1523/jneurosci.16-06-02027.1996

13. Katsimpardi L, Lledo PM. Regulation of neurogenesis in the adult and aging brain. Curr Opin Neurobiol. 2018;53:131–138. doi:10.1016/j.conb.2018.07.006

14. Soares R, Ribeiro F, Lourenço Di, et al. The neurosphere assay: An effective in vitro technique to study neural stem cells. Neural Regen Res. 2021;16(11):2229–2231. doi:10.4103/1673-5374.310678

15. Pilz GA, Bottes S, Betizeau M, et al. Live imaging of neurogenesis in the adult mouse hippocampus. Science (80- ). 2018;359(6376):658-662. doi:10.1126/science.aao5056

16. Wu Y, Bottes S, Fisch R, et al. Chronic in vivo imaging defines age-dependent alterations of neurogenesis in the mouse hippocampus. Nat Aging. 2023;3(4):380–390. doi:10.1038/s43587-023-00370-9

17. Murray KD, Liu XB, King AN, Luu JD, Cheng HJ. Age-related changes in synaptic plasticity associated with mossy fiber terminal integration during adult neurogenesis. eNeuro. 2020;7(3). doi:10.1523/ENEURO.0030-20.2020

18. Duan X, Chang JH, Ge S, et al. Disrupted-In-Schizophrenia 1 Regulates Integration of Newly Generated Neurons in the Adult Brain. Cell. 2007;130(6):1146–1158. doi:10.1016/j.cell.2007.07.010

19. Fernández-Hernández I, Rhiner C, Moreno E. Adult Neurogenesis in Drosophila. Cell Rep. 2013;3(6):1857–1865. doi:10.1016/j.celrep.2013.05.034

20. Ganz J, Brand M. Adult neurogenesis in fish. Cold Spring Harb Perspect Biol. 2016;8(7):1-21. doi:10.1101/cshperspect.a019018

21. Goldman SA, Nottebohm F. Neuronal production, migration, and differentiation in a vocal control nucleus of the adult female canary brain. Proc Natl Acad Sci U S A. 1983;80(8 I):2390-2394. doi:10.1073/pnas.80.8.2390

22. Eriksson PS, Perrfilieva E, BJörk-Eriksson T, et al. Erikkson Et Al (1998). Nat Med. 1998;4(11):1313-1317.

23. Rust R, Walker TL. Isolation and Culture of Adult Hippocampal Precursor Cells as Free-FloatingFree floatingNeurospheres. In: Deleyrolle LP, ed. Neural Progenitor Cells: Methods and Protocols. Springer US; 2022:33–44. doi:10.1007/978-1-0716-1783-0_3

24. Xiong F, Gao H, Zhen Y, et al. Optimal time for passaging neurospheres based on primary neural stem cell cultures. Cytotechnology. 2011;63(6):621–631. doi:10.1007/s10616-011-9379-0

25. Soares R, Ribeiro FF, Lourenço DM, et al. Isolation and Expansion of Neurospheres from Postnatal (P1&#8722;3) Mouse Neurogenic Niches. J Vis Exp. 2020;(159). doi:10.3791/60822

26. Walker TL, Kempermann G. One mouse, two cultures: Isolation and culture of adult neural stem cells from the two neurogenic zones of individual mice. J Vis Exp. 2014;(84):1–9. doi:10.3791/51225

27. Kalamakis G, Brüne D, Ravichandran S, et al. Quiescence Modulates Stem Cell Maintenance and Regenerative Capacity in the Aging Brain. Cell. 2019;176(6):1407–1419.e14. doi:10.1016/j.cell.2019.01.040

28. Urbán N, Blomfield IM, Guillemot F. Quiescence of Adult Mammalian Neural Stem Cells: A Highly Regulated Rest. Neuron. 2019;104(5):834–848. doi:10.1016/j.neuron.2019.09.026

29. Audesse AJ, Webb AE. Mechanisms of enhanced quiescence in neural stem cell aging. Mech Ageing Dev. 2020;191:1–20. doi:10.1016/j.mad.2020.111323

30. Pozzi D, Ban J, Iseppon F, Torre V. An improved method for growing neurons: Comparison with standard protocols. J Neurosci Methods. Published online 2017. doi:10.1016/j.jneumeth.2017.01.013

